# Global climate changes will lead to regionally divergent trajectories for ectomycorrhizal communities in North American Pinaceae forests

**DOI:** 10.1101/393009

**Authors:** Brian S. Steidinger, Jennifer M. Bhatnagar, Rytas Vilgalys, John W. Taylor, Thomas D. Bruns, Kabir G. Peay

## Abstract

Ectomycorrhizal fungi (ECMF) are partners in a globally distributed tree symbiosis that enhanced ecosystem carbon (C)-sequestration and storage. However, resilience of ECMF to future climates is uncertain. We sampled ECMF across a broad climatic gradient in North America, modeled climatic drivers of diversity and community composition, and then forecast ECMF response to climate changes over the next 50 years. We predict ECMF richness will decline over nearly half of North American Pinaceae forests, with median species losses as high as 21%. Mitigation of greenhouse gas emissions can reduce these declines, but not prevent them. Warming of forests along the boreal-temperate ecotone results in projected ECMF species loss and declines in the relative abundance of C demanding, long-distance foraging ECMF species, but warming of eastern temperate forests has the opposite effect. Sites with more ECMF species had higher activities of nitrogen-mineralizing enzymes, suggesting that ECMF species-losses will compromise their associated ecosystem functions.

## Introduction

Forecasting changes in the diversity and composition of microbial symbiont communities under anticipated future climates is valuable for concentrating conservation efforts(van der Linde *et al.* 2018) and predicting changes to ecosystem function (Bissett *et al.* 2013; Koide *et al.* 2014; Duffy *et al.* 2017). Loss of host species results in decreased ecosystem productivity and stability across a broad range of taxa (Duffy *et al.* 2017), including effects on microbes (Duffy *et al.* 2017; Laforest-Lapointe *et al.* 2017). Recent advances in the molecular methods of measuring species richness and composition have made it possible to characterize current continental scale diversity patterns of soil microbes (Tedersoo *et al.* 2012a; Talbot *et al.* 2014; Tedersoo *et al.* 2014; van der Linde *et al.* 2018). However, continental-scale forecasts under future climates are unavailable for most microbial guilds, making it difficult to predict the consequences of climate change to global biodiversity and ecosystem services. Here we predict how the species richness, relative abundance, and composition of ectomycorrhizal fungi (ECMF) in North American pine forests will change over the next 50 years.

ECMF are obligate plant symbionts that dominate global temperate and boreal forest soil communities and are implicated in most major ecosystem processes (Phillips *et al.* 2013). Two ECMF processes in particular may enhance how ecosystems will buffer the atmosphere against increased CO_2_ emissions: (1) the C-fertilization effect, where ECMF increase forest productivity and nutrient mobilization in response to increased atmospheric CO_2_ (Terrer *et al.* 2016); and (2) the Gadgil effect, where ECMF inhibit decomposition by free-living soil microbes (Gadgil & Gadgil 1971; Averill & Hawkes 2016). Because these functions enhance ecosystem C-sequestration and –retention, respectively, they have the potential to buffer the planet against climate change by reducing CO_2_-associated radiative forcing.

The ability of ECMF to acquire soil organic nitrogen (N) is hypothesized to mediate C-fertilization and the Gadgil effect. ECMF produce extracellular proteolytic and oxidative enzymes that liberate N from organic complexes. When ECMF transfer this N to their host trees under elevated CO_2_, it can fuel increased photosynthetic rates (Terrer *et al.* 2016). Similarly, by liberating N from organic complexes, ECMF are hypothesized to competitively inhibit free-living soil microbes that require N to decompose and respire soil organic matter (Averill *et al.* 2014). If EMCF-associated enzyme activity is dependent on the diversity and composition of ECMF communities (Talbot *et al.* 2013), then diversity losses and shifts in composition have the potential to compromise ECMF-associated C-sequestration and –retention.

Ecosystem services are also likely affected by shifts in the relative abundance of ECMF relative to other fungal guilds (inclusive of saprotrophs and pathogens) (Averill & Hawkes 2016), and also to shifts in the abundances of different nutrient foraging strategies within ECMF. Relative to short-distance hyphal exploration strategies, long distance foraging ECMF are associated with higher activities of organic N mineralizing enzymes (Hobbie & Agerer 2010; Tedersoo *et al.* 2012c) and potentially also relatively higher demands for host C (Agerer 2001; Deslippe *et al.* 2011; Fernandez *et al.* 2017). Thus, reducing the abundance of long-relative to short-distance foraging ECMF may result in declines in C-allocation belowground and the fungal N-mineralization hypothesized to drive the C-fertilization and Gadgil-effects.

Climate change can alter the community composition of ECMF by pushing fungi (Kipfer *et al.* 2010) or their host plants (Fernandez *et al.* 2017) outside their ranges of physiological tolerance (Pickles *et al.* 2012). However, most studies to date that have examined ECMF or whole fungal community responses to simulated climate change have found fairly small effects (Parrent *et al.* 2006; Tu *et al.* 2015; Fernandez *et al.* 2017; Mucha *et al.* 2018) relative to natural changes in fungal communities observed along large natural gradients of temperature and precipitation (Jarvis *et al.* 2013; Talbot *et al.* 2014; Tedersoo *et al.* 2014; Nottingham *et al.* 2016; Peay *et al.* 2017). Yet, few datasets currently exist with spatial resolution necessary to make accurate predictions of ECMF response to climate change across relevant geographic regions (Mohan *et al.* 2014).

To obtain the data needed to determine how ECMF communities are likely to change in altered climates, we used next-generation DNA sequencing to determine the species composition of ECMF in a series of 68 sites, each consisting of 26 soil samples taken with a consistent sampling design within a 40 x 40 m plot, spread across different EPA climatic regions in North America (Omernik & Griffith 2014) (Figure 1a). To isolate the effect of climate on ECMF communities while minimizing the known effects of vegetation biome type variation on microbial community structure and function (Fierer *et al.* 2012; Tedersoo *et al.* 2012b), we placed all of our sites in forests dominated by single species of tree from the family Pinaceae (an obligate ECMF host lineage).

**Figure 1.**
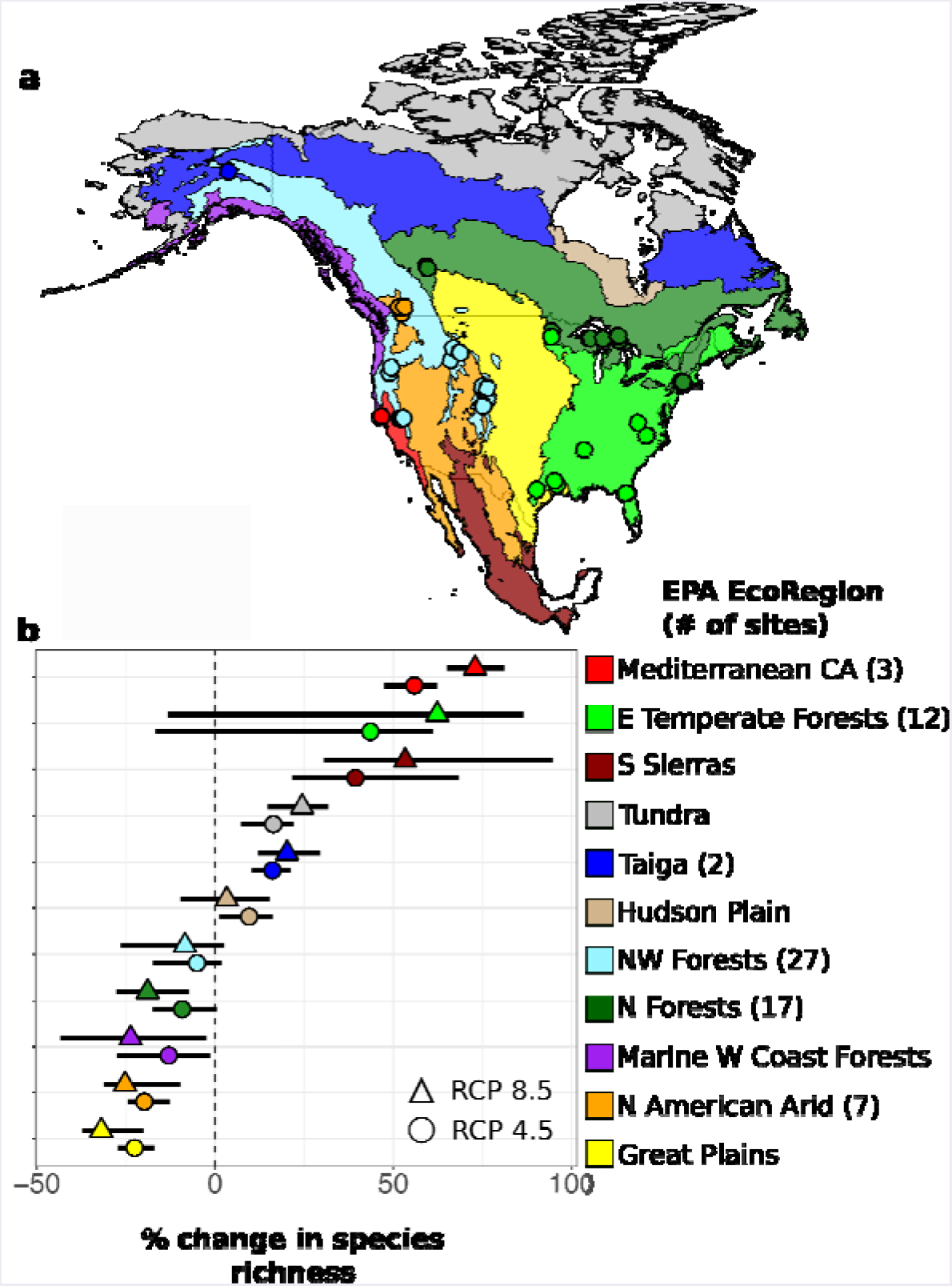
The predicted change in ECMF species richness by 2070 under two estimates of greenhouse gas emissions. (a) A map of North America shaded by EPA EcoRegion, with sites indicated by points. (b) Median predicted change in species richness by EcoRegion with error bars containing the inter-quartile range for RCP 8.5.

Recent historical climates (30-year means) influence soil nutrient availability and constrain the composition and function of ecosystems in ways that govern their response to changing climates (Karhu *et al.* 2014; Hawkes *et al.* 2017). To determine how historical climate (1960-1990) structures ECMF communities in our sites, we fit non-linear models to the relationships between ECMF species richness, relative abundance, and community dissimilarity and climate, in situ measurement of local soil chemistry, global rasters of soil chemical characteristics, and estimated total N-deposition. Using model selection criteria that maximize performance relative to model complexity, we eliminated all but climatic predictors from our models. Additionally, to identify ECMF community thresholds to changes along temperature gradients, we performed threshold indicator taxa analysis (Baker & King 2010). We then used our statistical models fit with 30-year climate means to forecast ECMF community changes under future (2070) climates, both with and without mitigations in greenhouse gas emissions. We inferred potential functional consequences of declines in ECMF diversity by comparing ECMF species richness to the activity of soil enzymes associated with hydrolysis and oxidation of organic substrates.

## Materials & Methods

### Sampling

We sampled 68 sites across North America by focusing on forests dominated by a single plant family, the Pinaceae (Table S1). The Pinaceae are ideal for exploring environment-community-function relationships across Kingdom Fungi because they have a broad distribution across North America and show low levels of host specificity for mycorrhizal fungi within the family (Rusca *et al.* 2006; Ishida *et al.* 2007). For example, North American pines readily associate with European ectomycorrhizal fungi (Vellinga *et al.* 2009) and co-occurring Pinaceae and angiosperms often share most common ectomycorrhizal fungi (Kennedy *et al.* 2003). Plots spread across North America were chosen with the help of local experts to find mature stands with high dominance of a single Pinaceae species (Fig. S1).

Sampling was carried out in 2011 and 2012 near the period of peak plant biomass production for a given region. In each plot, 13 soil cores were collected from a 40 m x 40 m grid (Fig. S1). To look at spatial turnover of community and function at the local to landscape scales, we ensured that each plot had at least one other plot located within a 1-50 km range (Fig. S1B). At each point in the plot, fresh litter was removed and a 14 cm deep, 7.6 cm diameter soil core was taken and immediately separated into a humic (O) horizon and mineral (A) horizon. This resulted in a total of 26 soil samples collected per plot (13 sample points x 2 horizons). After removal, soils were kept on ice until processed. Soils were sieved through a 2 mm mesh to remove roots and rocks and homogenized by hand. A ~0.15-0.25g subsample was placed directly into a bead tube from the Powersoil DNA Extraction Kit (MoBio, Carlsbad, CA USA), and the samples were stored at 4°C until DNA extraction. Before extraction, samples were homogenized for 30 seconds at 75% power using a Mini-Beadbeater (BioSpec, Bartlesville, OK USA). A second subset of soil core x horizon samples were stored in −80 C freezer (within 48 hours of collection) for soil chemical analysis.

### Soil Chemistry

Frozen soils were thawed and analyzed for pH in a 1:1 water ratio using a glass electrode. Total extractable ammonium and nitrate concentrations were analyzed in 2.0 M potassium chloride extracts of each soil sample using a WestCo SmartChem 200 discrete analyzer at Stanford University. For site-level values, we took the average of all soil cores processed for each site. Soil chemical variables were included in statistical models but later dropped during model selection (see Statistical Analysis and Supplemental Materials).

### Enzyme Assays

We assayed the potential activities of six extracellular enzymes involved in soil carbon and nutrient cycling (using methods described in (Talbot *et al.* 2014): B-glucosidase (BG, which hydrolyzes cellobiose into glucose), polyphenol oxidase (PPO, which oxidizes phenols), peroxidase (PER, including oxidases that degrade lignin), acid phosphatase (AP, which releases inorganic phosphate from organic matter), N-acetylglucosaminidase (NAG, which breaks down chitin), and leucin-aminopeptidase (LAP, which breaks down polypeptides). Potential enzyme activities in bulk soil were measured separately for individual organic and mineral horizon samples using fluorometric and colorimetric procedures (German *et al.* 2011) on a microplate reader (n=253).

### Molecular Methods

To characterize fungal communities we sequenced the internal transcribed spacer (ITS) region of the nuclear ribosomal RNA genes, the official barcode of life for fungi (Schoch *et al.* 2012). Because of improvements in technology during the course of this project, soil samples were sequenced using two different platforms. Soil samples from twenty-five sites were sequenced via 454 pyrosequencing as per Talbot *et al.* 2014. The remaining 43 sites were sequenced on an Illumina MiSeq at the Stanford Functional Genomics Facility using the primer constructs and protocols from (Smith & Peay 2014). Using a common set of soil samples from this study sequenced on both platforms we have previously demonstrated that both richness and species composition are highly reproducible and strongly correlated between the two platforms (Smith & Peay 2014) so that combining samples should not cause any bias in our analyses.

### Bioinformatics

Samples sequenced on the 454 platform were cleaned and denoised in QIIME (Caporaso *et al.* 2010) (Reeder & Knight 2010), after which we extracted the ITS1 region (Nilsson *et al.* 2010). For samples sequenced on the Illumina platform, samples were first trimmed using Cutadapt (Martin 2011) and Trimmomatic (Bolger *et al.* 2014), and then merged using USEARCH (Edgar 2010). At this point cleaned 454 and Illumina sequences were merged into a single FASTA file, where sequences were dereplicated, clustered into species level operational taxonomic units (OTUs) at 97% sequence similarity using USEARCH. We removed all singletons and chimeras, and dropped occurrences <0.025% of relative sequence abundance within a sample to account for tag-swapping (Carlsen *et al.* 2012). We took a two-step approach to assigning taxonomic identity, first using the BLAST tool with the UNITE reference database (Koljalg *et al.* 2013) to eliminate potentially non-fungal taxa; and next using the naïve Bayesian classifier from the Ribosomal Database Project (RDP) (Wang *et al.* 2007) along with the Warcup ITS reference set (Deshpande *et al.* 2016) to assign taxonomy. OTUs with sufficiently confident taxonomic assignments were then matched to functional guilds using the FUNGUILD database (Nguyen *et al.* 2016). The relative abundance of ECMF exploration strategies (% of ECMF OTUs) were assigned by matching ECMF genera against published lists (Agerer 2006; Tedersoo & Smith 2013) , which are available online (www.deemy.de). Strategies were assigned to the following categories according to (Fernandez *et al.* 2017): contact short (CS), contact medium (CM), and medium long (ML). Full description of the bioinformatic methods are available in the online supplement.

### Statistical Analyses

To correct for variability in DNA sequencing depth between samples from the two sequencing platforms, we rarefied our sequences to an even-depth of 17,273 sequences. The even-depth was determined after aggregating samples to their respective sites. To determine the relative abundance of ECMF sequences relative to all fungal sequences, we also summed the total number of ECMF sequences and divided it by the sum of all fungal sequences.

Recent-historical climates (30 year means) are commonly used to model the bioclimatic variables that shape current species distributions and responses to climate change (Hijmans & Graham 2006). To determine the role of climate in shaping the current and future distributions of ECMF species richness and community composition, we downloaded the following 30 year mean (1960-1990) and projected future (2070) bioclimatic variables from World Clim version 1.4 (Hijmans *et al.* 2005): mean annual temperature (°C), temperature seasonality (standard deviation of monthly temperatures), mean annual precipitation (mm), and seasonality of precipitation (coefficient of variation in monthly precipitation). These variables capture both the range and central tendency of climate factors that are demonstrated to affect fungal communities and exhibit low co-linearity (variance inflation factor < 3), which minimizes spurious fits between predictor and response variables. We extracted data for our 68 sites from these climate rasters.

We fit generalized additive models (GAMs) of the ECMF species richness, relative abundance (out of all fungi), and relative abundance of CS, CM, and ML exploration strategies of each site as a function of the bioclimatic variables using the mgcv package in R (Figure S7). We choose to use non-parametric GAMs, rather than linear models, so that our statistical models would have sufficient flexibility to capture curvilinear responses of EMCF diversity and abundance to environmental factors. However, a potential drawback of this approach is the potential to over-fit data, predicting environmental responses that are difficult to interpret biologically. In order to navigate these twin pitfalls, we constrained our model fits to four knots to allow for threshold, saturating, uni- and bi-modal responses to environmental variables.

In addition to the four climatic variables for which we have recent-historical and projected 2070 data, we also considered GAMs with only *in situ* soil predictors (soil pH, NH4-N), only soil predictors from global rasters of surface horizon chemistry [soil pH in KCl (Hengl *et al.* 2017) and total N density (Task 2000)], only estimates of total atmospheric N-deposition (Hember 2018), and combinations of climate with soil *in situ*, soil raster, and N-deposition predictors. We selected our climate-only model based on its superior performance relative to model-complexity according to the generalized cross validation (GCV) statistic (full details in Supplementary Materials). In contrast with soil data, climate variables are also available across North American pine forests for both recent-historical and projected future scenarios (contingent on anthropogenic greenhouse emissions), allowing us to predict and project ECMF species richness, abundance, and community composition across time and space.

We used our statistical models to generate predictions over North America, under both historical and future climate conditions. We constrained our predictions to the spatial extent of the distributions of the twelve most abundant pine tree species from our plots. Tree species distributions were derived from the United States Geological Survey (USGS) shapefiles, which we dissolved into one composite range using the “raster” package in R. To summarize our results by region, we subset our predictions according to EPA vegetation zones (Omernik & Griffith 2014) . In Figure 1ab, the EcoRegions temperate sierras, semi arid highlands, and tropical dry forests were combined into the composite Ecoregion “S. Sierras.” This was done due to the similar climate of the small areas of the constituent regions that overlap with the distribution of pine-forests.

To project our statistical models to future climates (the year 2070), we downloaded predicted climate rasters from 17 different global climate models (GCMs) that have been incorporated into the Coupled Model Intercomparison Project Phase 5 (CMIP5) (Table S5). While these 17 GCMs differ in complexity, each simulates anthropogenic changes using two greenhouse emission scenarios, corresponding to Relative Concentration Pathways (RCP) of 4.5 and 8.5 (Allen *et al.* 2014). In order to offset prediction-errors associated with individual GCMs (e.g., (Pierce *et al.* 2009), we projected ECMF species richness, relative abundances, and composition using a consensus-GCM, which is the average predicted climate for each pixel across all 17 GCMs. To plot the percent change in species richness, we took the predicted [(future richness - historical richness)/(historical richness)].

In order to analyze ECMF community composition, we first transformed our OTU table into a table of the proportion of sequences represented by each species at our sites. We then derived a Bray-Curtis dissimilarity matrix for each pair of sites. The relationship between geographic distance and difference in climate predictors of each site was analyzed using generalized dissimilarity models (GDM) using the package “gdm” in R (Manion *et al.*). We used the fitted GDM model to project the change in dissimilarity among forest sites in historical vs. projected climates. For the future climate, we scaled the first PC axes of predicted community composition from 0-255 and plotted the points using three dimensional color scaling [PC A 1 (green), PCA 2 (red), PCA 3 (blue)].

In order to identify responses of individual ECMF taxa to changes in temperature gradients, we performed Threshold Indicator Taxa ANalysis (TITAN) using the “TITAN2” package in R (Baker & King 2010). First, we aggregated our OTU table to the level of assigned species name and removed all species that occurred in fewer than 4 sites. Next, we used TITAN to find the individual ECMF species responses to gradients of mean annual temperature, which our analyses identified as having two different ECMF-diversity optima. The analysis returns two metrics for each species: (1) a change point, which splits each species’ abundances into two classes along a environmental gradient (above and below a set temperature) in a way that maximizes the fidelity of species’ association for one of the two classes; ands (2) a standardized z-score, where the magnitude is proportional to the sensitivity of the species to change and the sign indicates that the species’ increases or decreases in relative abundance when temperatures exceed the change point (positive and negative z-scores, respectively). For plotting purposes, we display only pure and reliable indicator taxa [e.g., (van der Linde *et al.* 2018)], We identified the temperature thresholds for ECMF communities as the peaks in the cumulative distributions of the negative and positive-z values, respectively.

The mean activity of each enzyme in all cores (organic and mineral horizons) were calculated for each site and log-transformed. Because activity of individual enzymes were often correlated we express enzyme activity in terms of the first two axes from principal component analyses of activities across the 25 sites where enzyme activity was measured.

## Results

Climate variables explained 58% of the deviance in ECM species richness (range 33-199 species per plot), 41% of the deviance in ECMF relative abundance (7-78% of sequences), and 41% of species composition (0.6-1 dissimilarity) among sites (Table 1). The most species-rich and relatively abundant ECMF communities are associated with sites with high seasonality in temperature and precipitation (Figure 2ab). Seasonality also explains the most variability in ECMF species composition, with the largest differences associated with differences in temperature seasonality (Figure 2c).

**Figure 2.**
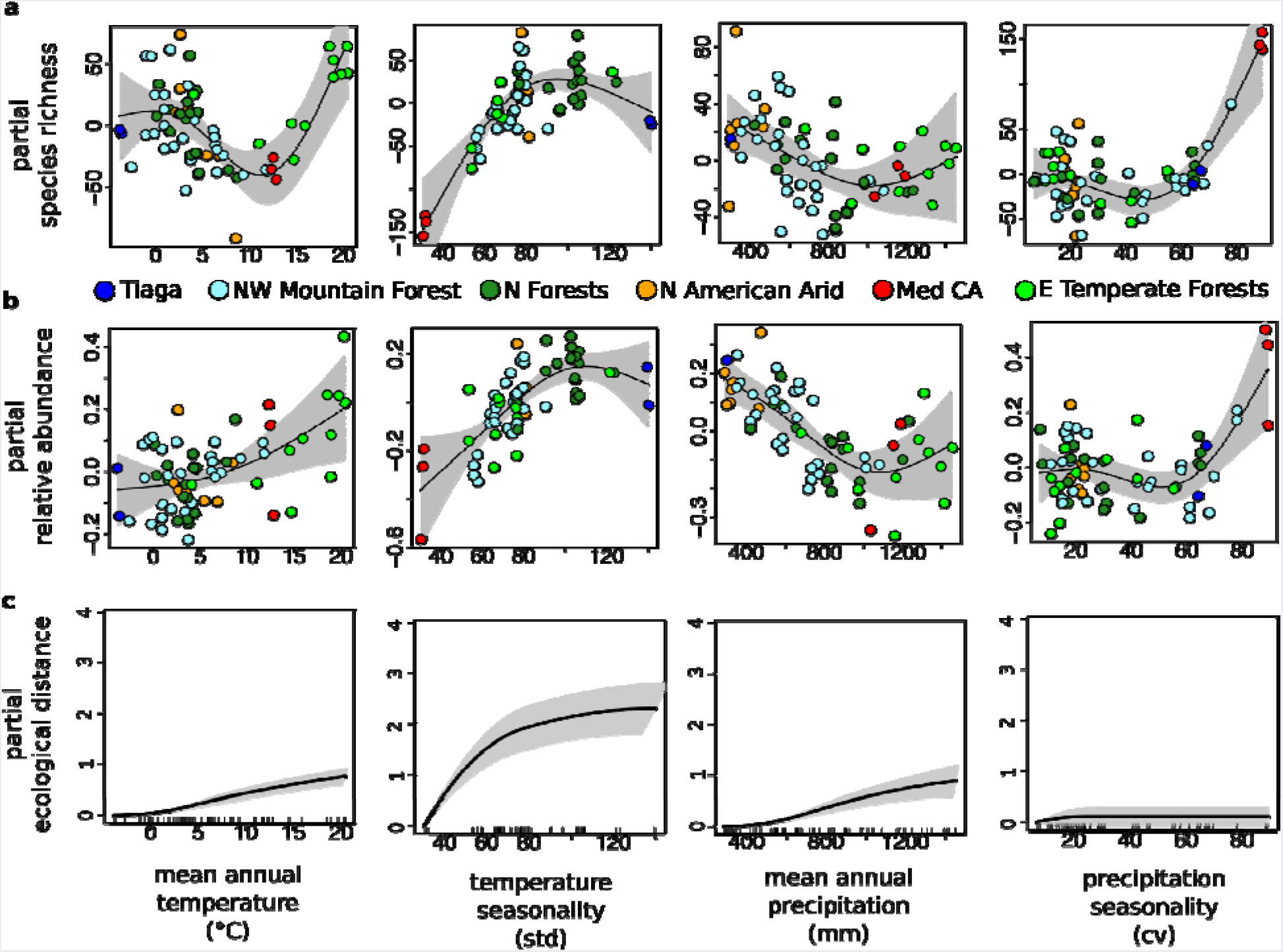
The predictions of partial regression and best-fit splines for general-additive models for species richness (a) and relative abundance (b) by historical (1960-1990) climate, with shading along the 95% confidence interval. Total species richness and relative abundance for each site are equal to the sum of the four predictions and an intercept value (97.04 and 0.48 for species richness and relative abundance, respectively). Points are colored according to EPA EcoRegion as in Figure 1. (c) The expected difference in ECMF community composition, measured as the partial ecological distance, among sites according to generalized dissimilarity models, with lines on the x-axis indicated empirical values, while shading represents the 95% confidence interval after sampling 70% of sites 10 times.

**Table 1.** Current summary statistics for (a) ECMF species richness, (b) relative abundance, and (c) Bray-Curtis dissimilarity, along with significance of climate variables in general-additive models [(k)nots=4] for (a) and (b) and a summary of the general-dissimilarity model for (c, k=3).

| **(a) species richness (mean = 97.4, median = 98.5, iqr = 59.5)** |  |  |  |  |
| --- | --- | --- | --- | --- |
| <u>climate variable</u> | <u>Estimated</u><br><u>degrees of</u><br><u>freedom (df)</u> | <u>Ref.df</u> | <u>F-value</u> | <u>p-value</u> |
| Mean annual temperature | 2.89 | 2.98 | 9.98 | <0.001 *** |
| Temperature seasonality | 2.59 | 2.89 | 10.64 | <0.001*** |
| Mean annual precipitation | 2.15 | 2.54 | 5.45 | 0.01** |
| Precipitation seasonality | 3.00 | 3.00 | 7.59 | <0.001*** |
| Deviance explained = 58.3%; GCV = 845.43; Scale est. = 700.78 |  |  |  |  |
| **(b) relative abundance (mean = 0.48, median = 0.52, iqr = 0.17)** |  |  |  |  |
| Mean annual temperature | 1.75 | 2.07 | 11.23 | 0.07 . |
| Temperature seasonality | 2.48 | 2.80 | 13.54 | <0.001*** |
| Mean annual precipitation | 1.75 | 2.07 | 11.23 | <0.001*** |
| Precipitation seasonality | 2.81 | 2.96 | 4.52 | <.01** |
| Deviance explained = 54.3%, GCV = 0.014, Scale est. = 0.012 |  |  |  |  |
| **(c) Bray-Curtis dissimilarity (mean = 0.94, median = 0.97, iqr= 0.07)** |  |  |  |  |
| Null Deviance = 154. 11; GDM Deviance = 90.6; Deviance explained = 41.2% |  |  |  |  |

ECMF respond to historical climates differently by region. The most species-rich ECMF communities in the northern and northwest mountain forests are cold and dry (0°C and < 1 m mean annual temperature and precipitation, respectively). By contrast, the most species-rich ECMF communities in eastern temperate forests are hot and wet (> 12°C, >1 m precipitation, figure 2a). Using general-additive models, which fit continuous, curvilinear responses along temperature gradients, we found a bimodal relationship between species richness and mean temperature, with separate cold- and hot-diversity optima (Figure 2a). As a result, our models predict that warming decreases species richness in the relatively cold north/northwest forests and increases species richness in the eastern temperate forests.

Relative abundance and species richness of ECMF are correlated (R^2^=0.23, p<0.01) and respond similarly to variability in climate (Figure 2ab). However, unlike ECMF species richness, ECMF relative abundance increases with temperature even in forests with mean annual temperatures < 12°C. As a result, our statistical models predict partial increases in ECMF relative abundance with warming, although the net-effect is contingent on climate changes to precipitation and seasonality (Figure 2b).

Similar to our models of ECMF species richness, the relative abundance of long-distance foraging strategies increases with mean temperature in eastern temperate forests with mean annual temperatures > 12°C and declines with mean temperature in southern boreal forests (Figure S7c), By contrast, both short and medium-distance foragers increase in relative abundance with temperature increases in the colder north / northwestern EcoRegions (Figure S7ab). As a result, our models predict that sites that lose species with increased temperatures should also decline in the abundance of long-distance foraging strategies.

Threshold indicator taxa analysis identifies ECMF species that reliably respond either negatively or positively to increases in mean annual temperature (Figure 3a). Based on the peaks of the cumulative distributions of the change points for ECMF species with negative and positive temperature associations (Figure 3b), we identify two ECMF-community thresholds: (1) a cold threshold at 3 °C, which is associated with declines among threshold indicator species with negative z-scores; and (2), a hot threshold at 12 °C, which is associated with increases among threshold indicator species with positive z-scores. Notably, these ECMF community thresholds occur near the inflection points for the response of overall ECMF species richness to mean annual temperature (Figure 1a).

**Figure 3.**
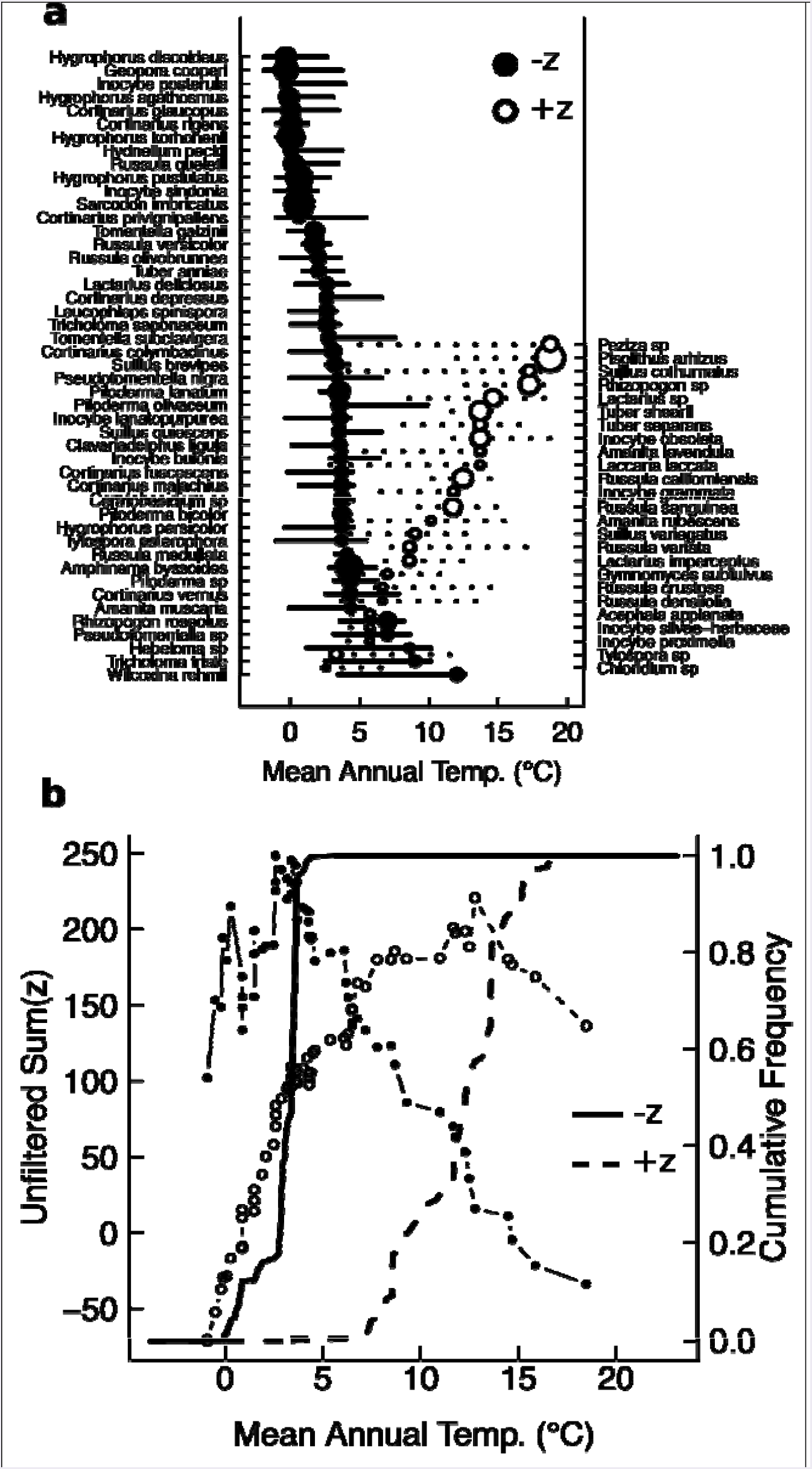
(a) The change points (circles) and 95% confidence intervals for ECMF species with negative and positive responses to increasing mean annual temperature (-z and +z, respectively). (b) The sum of z-scores (lines with points, left axis) and cumulative distribuiton of change points (right axis) for ECMF species with negative and positive z-scores. Peaks in the unfiltered sum(z) and sharp increases in cumulative frequency indicate ECMF community thresholds for change along mean annual temperature gradients.

We compared model predictions using 30-year mean climates (1960-1990) to climate projections for 2070 both with and without mitigation in greenhouse gas emissions (RCP 4.5 and 8.5, respectively). Our models predict that both ECMF diversity and the abundance of longdistance foragers decline at the temperate-boreal ecotone, which includes the southern extent of the northern forests, the northwest mountain forests, and the marine west coast forests (Figures 1b,3, 6). By contrast, our models predict increases in ECMF species richness in south/southeastern forests, including the southern extent of the eastern temperate and temperate sierran forests of Northern Mexico (Figure 1a, Figure 4a). The projected shifts in community composition, which are independent from projected changes in species richness, are also more severe at the southern and western extent of the northern and northwest mountain forests and along the eastern temperate forests (Figure 4c). With mitigation in greenhouse gas emissions, median loss of species richness and the geographic extent of those declines shrink from 21 to 14% and from 48 to 44% of pine forests, respectively. Species gains are also more extensive and intense with higher greenhouse-gas emissions (Figures 1b,4).

**Figure 4.**
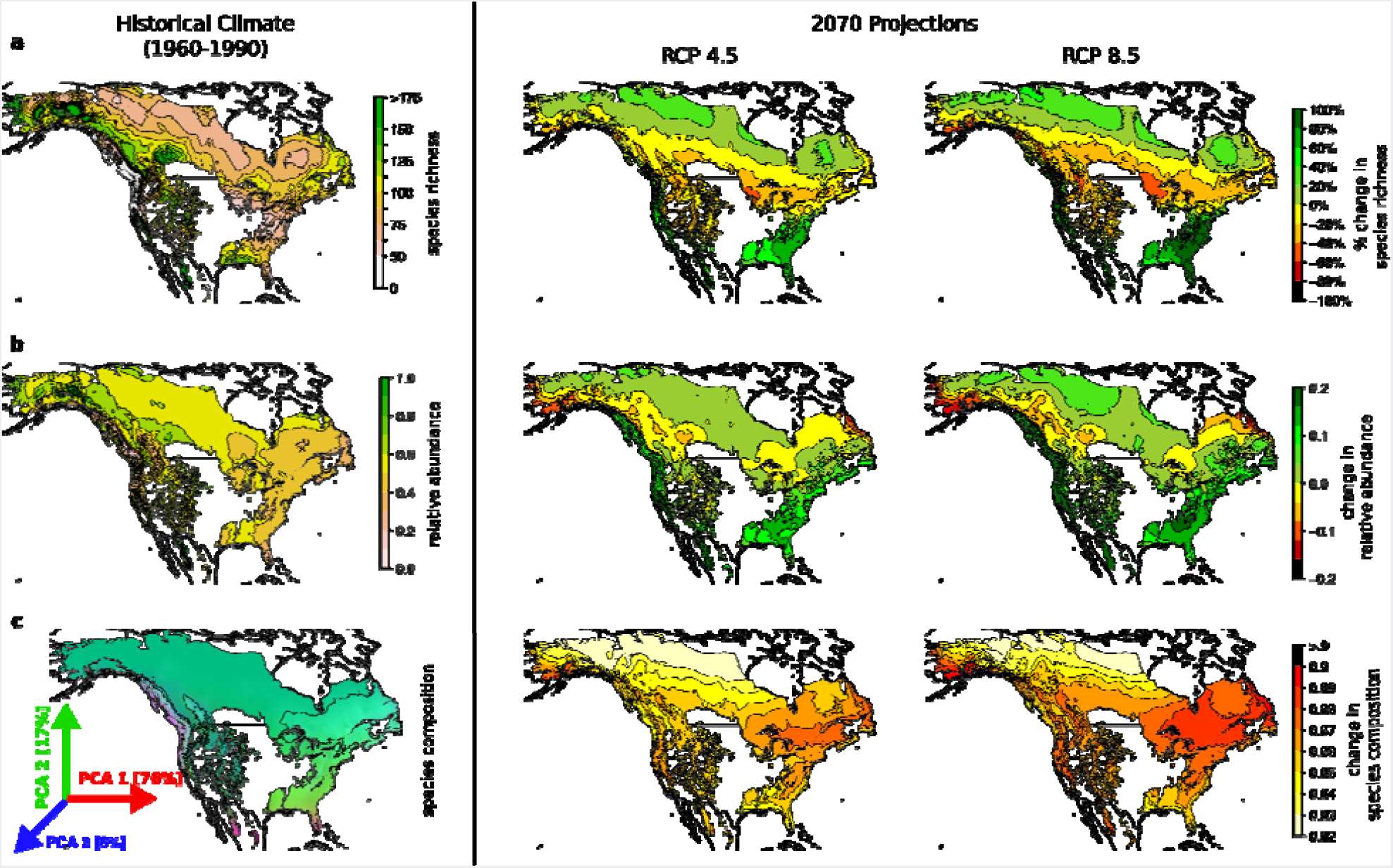
Predicted maps (right) using historical climate and general-additive models for (a) species richness, (b) relative abundance and the (c) first three PCA axes indicated by green, red, and blue color scaling using a general dissimilarity model for species composition [with % of predicted variance explained]. Contour lines for species composition (c) delineate four clusters in the three dimensional PCA space. The projected changes for 2070 under different RCP scenarios are plotted to the left. Regions of high ECMF predicted species richness (a) at the temperate-boreal ecotone and throughout northwest mountain and marine west coast forests are particularly vulnerable to species loss, while eastern temperate forests are expected to increase to species richness.

Sites differed in the soil activities of five different extra-cellular enzymes, which we represent with the first two axes of principal component analysis (explaining 56 and 21% of enzymatic variability, Figure 6a). All assayed enzymes load positively onto PC axis 1, while axis 2 discriminates between oxidative (positive loading) and hydrolytic enzymes (negative loading, with the exception of leucine-aminopeptidase). Both total and oxidative enzyme activity are correlated with ECMF species richness using single regression (Figure 6bc), but the relationship is statistically significant only for oxidative enzymes (R^2^=0.17, p=0.04).

**Figure 6.**
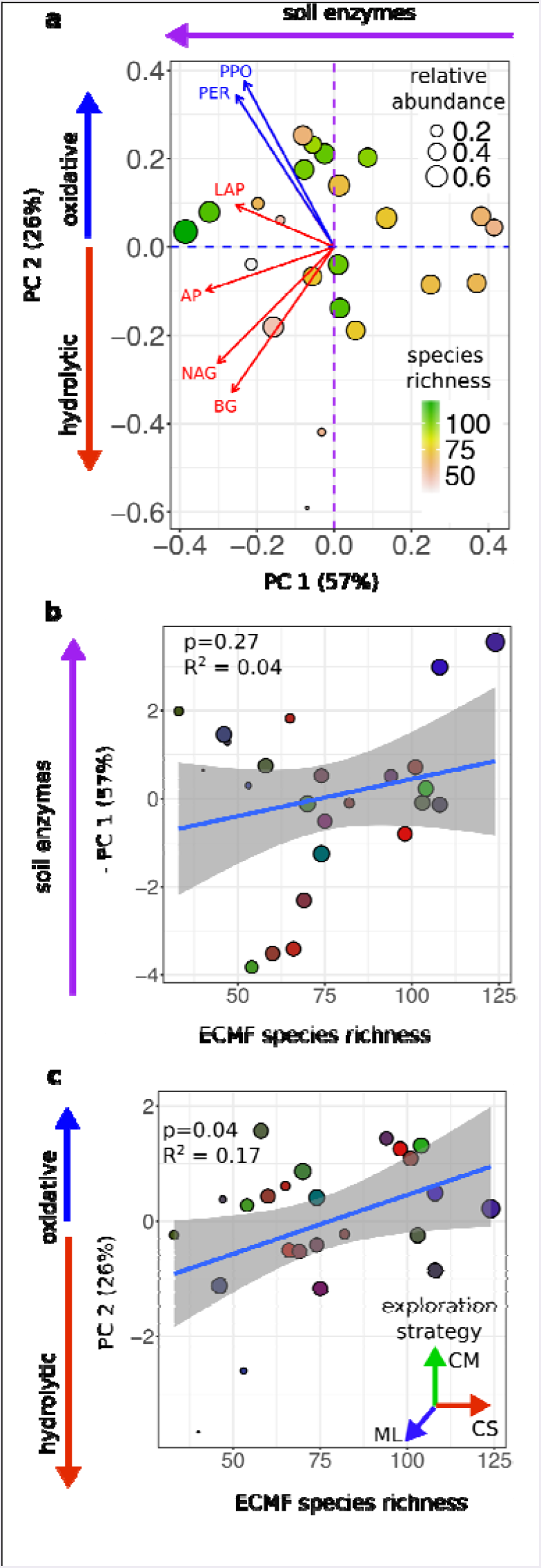
(a) Plot of forest sites against the first two axes of principal component analysis for logtransformed enzyme activity. Blue vector indicate the loading values of oxidative enzymes (PPO; PER, peroxidase), while red vectors indicate loading values of hydrolytic enzymes (LAP, leucinaminopeptidase; AP, acid phosphatase; NAG, Nacetylegulocsaminidase; BG, B-glucosidase). All enzymes load negatively onto PC 1, while PC 2 discriminates between oxidative and hydrolytic enzyme activity. (b) and (c) show the correlation between ECMF species richness and -PC 1 (notsignificant) and PC 2 (significant). All points are sized according to the relative abundance of ECMF relative to all fungi (legend in panel a), while points in (b) and (c) are colored using 3D color scaling of the relative abundance of different hyphal exploration strategies (CS, contact-short; CM, contact-medium; ML, medium-long).

## Discussion

With no mitigations in greenhouse gas emissions, we predict that as many as 48% of North American pine forests will lose 21% of their ECMF species due to climate changes in the next 50 years. Predicted declines in ECMF diversity are associated with the temperate-boreal ecotone, which is already vulnerable to global change due to warming-associated declines in growth and photosynthetic rates of southern boreal trees (Reich & Oleksyn 2008). Our predictions are generated using a model of historical climate on current ECMF diversity and composition that yields two key findings: (1) continental diversity patterns of ECMF in Pinaceae forests are driven primarily by differences in temperature and precipitation seasonality and (2) the ECMF diversity of a forest can either increase or decrease with mean annual temperature, contingent on its association with cold- or hot-diversity optima.

Our models explain a substantial fraction of variability in ECMF diversity and community structure using climate only, with soil N-availability and anthropogenic N-deposition being dropped as predictors during model selection due to their low predictive power relative to model-complexity (Supplementary Materials). This negative result with respect to N-deposition contrasts with some regional studies of ECMF (Pardo *et al.* 2011; Jarvis *et al.* 2013; Suz *et al.* 2014; Batstone *et al.* 2017), including a recent continental-scale analysis of ECMF community composition across western Europe (van der Linde *et al.* 2018). By contrast, we found that once the effects of climate gradients were taken into account, our multiple regression models fit small and statistically insignificant responses of ECMF diversity to total N deposition (Figure S6). While we acknowledge that there are many potential drivers of ECMF community structure, our study design, which focuses exclusively on ECMF in Pinaceae dominated forest stands, allowed us to isolate the large and regionally divergent responses of ECMF communities to spatial-temporal climate gradients.

The most diverse EMCF forests in our network had highly seasonal temperature and precipitation. Seasonal forests are also associated with ephemeral flushes of nutrients (Voříšková *et al.* 2014), which ECMF can rapidly absorb, store in networks of soil mycelia, and transfer to tree hosts at later times (Read 1991). The higher predicted ECMF diversity associated with seasonality in precipitation mirror results from experimental manipulations on fungal communities (Hawkes *et al.* 2011) and suggests ECMF diversity may be associated with a storage effect of seasonal specialists (Chesson 2000).

The regionally opposite effects of temperature gradients we observed for ECMF diversity and composition are consistent with regionally opposite effects of climate on host-tree physiology. We found that ECMF diversity and long-distance forager abundance decline with increasing mean temperatures in sites from northern and northwest mountain forests, but increase with temperature in sites from eastern temperate forests. As a result, our models predict that ECMF diversity and long-distance forager abundance will decrease in sites along the temperate-boreal ecotone, but increase in eastern temperate forests. Warming of boreal tree species growing near the boreal-temperate ecotone has also been shown to reduce tree growth and photosynthetic rate (Reich & Oleksyn 2008; Reich *et al.* 2015; Fernandez *et al.* 2017), which causes trees to allocate less C to ECMF (Fernandez *et al.* 2017). By contrast, stimulated warming does not result in declines in photosynthetic rates for temperate tree species, which are adapted to warmer climates, or in boreal tree species growing at their colder, northern range limits (Reich & Oleksyn 2008). Thus, the qualitatively different effects of climate we detected on ECMF species richness and composition could reflect the climate-envelopes of their associated host trees, with projected declines in species richness occurring only in regions where ECMF host tree performance declines with increasing temperature.

Additionally, adaptation of ECMF species to different climate envelopes can explain qualitatively different community responses to temperature and precipitation (Lehto *et al.* 2008; Malcolm *et al.* 2009) (Hawkes & Keitt 2015). ECMF communities associated with different sites have completely separate species compositions (mean dissimilarity of 0.97 out of 1, 42% species endemic to a single site), which is consistent with the generally high spatial turnover among soil fungal communities (Talbot *et al.* 2014). The most dissimilar ECMF communities also have the most contrasting climates (e.g., northern forests with hot summers and cold winters vs. pacific coastal and southeastern temperate forests that lack seasonality, Figure 2c), such that different ECMF species are associated with the cold- and hot-diversity optima. Similarly, threshold indicator taxa analysis along mean annual temperature gradients identify ECMF community thresholds at ~3°C (Figure 3b), such that a host of cold-adapted ECMF species should decline with increasing temperature (Figure 3a). However, whereas small-scale studies of stimulated warming generally find low or absent effects on ECMF diversity (Parrent *et al.* 2006; Tu *et al.* 2015; Fernandez *et al.* 2017; Mucha *et al.* 2018), our models predict that addition to EMCF compositional changes, larger scale climate alteration will result in substantial diversity losses and gains across North America.

The magnitude of ECMF species losses and gains are contingent on the decisions of human policymakers, though even with mitigation of greenhouse gas emissions the outcomes are not qualitatively different. If greenhouse gas emissions are capped by 2040, median loss of species richness and the geographic extent of those declines shrink from 21 to 14% and from 48 to 44% of pine forests, respectively. Conversely, the predicted increase in ECMF diversity and abundance have the potential to increase C sequestration and retention in both far northern and south/southeastern forests. These increases are also more extensive and intense with higher greenhouse-gas emissions (Figures 1b, 3,4). However, while ECMF differ in foraging and dispersal strategy and enzymatic abilities, which can lead to positive diversity-seedling growth relationship from a range of 1-4 ECMF species in experimental tree seedlings (Baxter & Dighton 2001, 2005), it remains unclear how a system with ~100 ECMF species will respond to a loss or gain of 25-30%. Despite uncertainty of the magnitude of this effect, recent work suggests that species loss results in declines in ecosystem productivity and resiliency across a broad range of taxa (Duffy *et al.* 2017), including plant-microbial symbionts (Laforest-Lapointe *et al.* 2017).

The most likely functional fallouts for forest ecosystems predicted to lose ECMF diversity, such as northern and northwest mountain forests, are decreased C sequestration (via declining productivity (Terrer *et al.* 2016)) and decreased C-retention (via relaxed inhibition of free-living microbes (Averill & Hawkes 2016)). Two lines of evidence from our data support this functional shift: (1) forests projected to loss ECMF species in altered climates are also projected to decline in the relative abundance of C-demanding, long- and medium-distance exploration strategies (Figure 5abc); and (2) sites with higher ECMF species richness have higher activity of oxidative enzymes associated with N mineralization and the slow-decomposition of soil C (Figure 6). Thus, the loss of long- and medium-distance foraging strategies could result in less fixed-C being sequestered below-ground, while decreased enzymatic function associated with declines in ECMF diversity could compromise the ECMF-associated C-fertilization and Gadgil effects in altered climates (Averill *et al.* 2014; Averill & Hawkes 2016).

**Figure 5.**
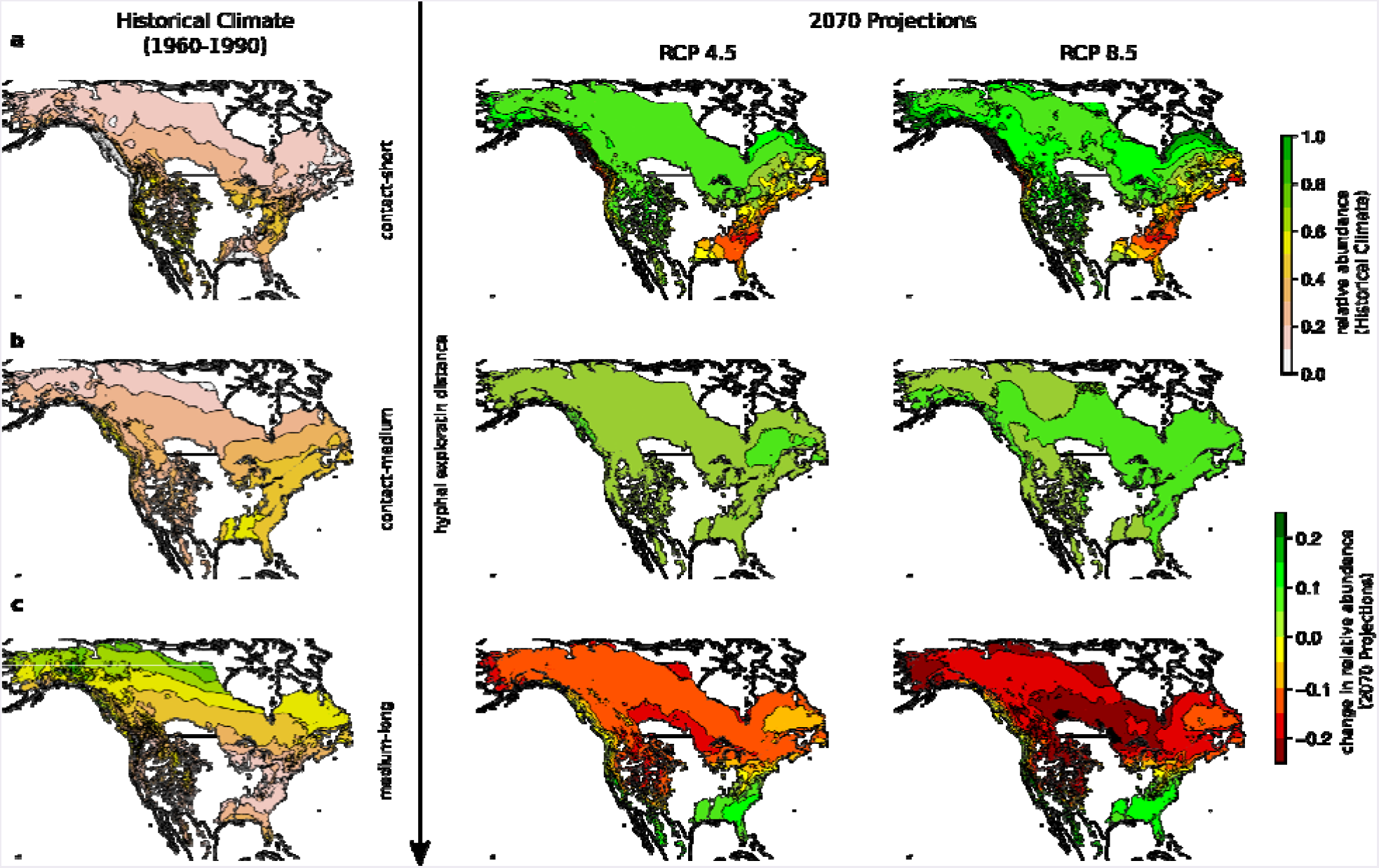
Predicted relative abundance (% of ECMF sequences) of (a) contact-short, (b) contact-medium, and (c) medium long hyphal exploration strategies under historical climates (right, top key) and projected changes in relative abundance under two different C emissions scenarios for 2070 (left, bottom key). Contact-short strategies increase sharply in northern / northwest forests and decrease in southeast temperate forests. Medium-long strategies have the opposite pattern.

Forecasting continental changes in ECMF communities is a first-step in projecting the functional consequences of those changes. In summary, over the next 50 years we predict climate
change will cause ECMF species richness to contract by 16-22% in north/northwestern pine forests of North America and expand by 21-28% in south/southeast pine forests. These changes have the potential to expand and contract the role ECMF play as boosters of forest productivity and inhibitors of soil respiration by free-living microbes.

## Data Availability

Upon acceptance for publication, data will be archived in Dryad.

